# Maternal gut microbiota *Bifidobacterium* promotes placental morphogenesis, nutrient transport and fetal growth in mice

**DOI:** 10.1101/2021.07.23.453407

**Authors:** Jorge Lopez-Tello, Zoe Schofield, Raymond Kiu, Matthew J. Dalby, Douwe van Sinderen, Gwénaëlle Le Gall, Amanda N Sferruzzi-Perri, Lindsay J Hall

**Author notes:** **Corresponding author names:** Jorge Lopez-Tello, Amanda N Sferruzzi-Perrri, Lindsay J Hall ( I). Contributed equally (Jorge Lopez-Tello, Zoe Schofield, Amanda N Sferruzzi-Perri, Lindsay J Hall).

## Abstract

The gut microbiota plays a central role in regulating host metabolism. While substantial progress has been made in discerning how the microbiota influences host functions post birth and beyond, little is known about how key members of the maternal gut microbiota can influence feto-placental growth. Notably, in pregnant women, *Bifidobacterium* represents a key beneficial microbiota genus, with levels observed to increase across pregnancy. Here, using germ-free and specific-pathogen-free mice, we demonstrate that the bacterium *Bifidobacterium breve* UCC2003 modulates maternal body adaptations, placental structure and nutrient transporter capacity, with implications for fetal metabolism and growth. Maternal and placental metabolome were affected by maternal gut microbiota (*i*.*e*. acetate, formate and carnitine). Histological analysis of the placenta confirmed that *Bifidobacterium* modifies placental structure via changes in *Igf2P0, Dlk1, Mapk1* and *Mapk14* expression. Additionally, *B. breve* UCC2003, acting through *Slc2a1* and *Fatp1-4* transporters, was shown to restore fetal glycaemia and fetal growth in association with changes in the fetal hepatic transcriptome. Our work emphasizes the importance of the maternal gut microbiota on feto-placental development and sets a foundation for future research towards the use of probiotics during pregnancy.

## Introduction

All nutrients and metabolites required for feto-placental growth are provided by the mother, which in turn is thought to be influenced by the maternal gut microbiota through breakdown of complex dietary components [1]. During gestation, liberated metabolites may be used by the placenta for morphogenesis, and transported across the placenta for use by the fetus for growth and development [2, 3]. This is highly important across gestation, particularly at later stages, when fetal growth is maximal. Notably, there are also alterations in the maternal microbiota throughout pregnancy with levels of the bacterial genus *Bifidobacterium* rising from trimester 1 onwards [4–6]. Failure of the mother to provide nutrients and metabolites to the fetus can result in pregnancy complications including small for gestational age, fetal loss and stillbirth. However, the contribution of the maternal gut microbiota in determining fetal outcomes is largely unexplored. Knowledge in this area would be highly valuable for developing treatments to improve fetal growth, with benefits for population health.

Studies performed with germ free (GF) mice have identified that the microbiota is a key regulator for adequate development, early immune education and metabolism [7–11]. However, little is known about how maternal gut microbiota influences feto-placental growth and placental structure and function. Here, we hypothesized that the maternal gut microbiota, and specific microbiota members, regulate fetal growth by modulating placental development and nutrient supply. We tested this hypothesis by comparing conceptus growth across a range of microbiome complexity; using conventional specific-pathogen-free (SPF) mice as a model for standard microbial colonization, and as a base line to define correct feto-placental growth [11]; GF mice which represent a completely clean and naïve microbiome system; and a mono-colonized maternal GF model – GF mice colonized with *Bifidobacterium breve* UCC2003 (group referred throughout the manuscript as BIF) [12]. *Bifidobacterium*, including *B. breve* UCC2003, is known to beneficially modulate the wider gut microbiota and host responses [13–15]. It is defined as a probiotic “live microorganisms, which when ingested or locally applied in sufficient numbers confer one or more specified demonstrated health benefits for the host” (FAO/WHO; [16]). Therefore, *B. breve* may represent a suitable option for treating pregnancy complications by exerting metabolic effects on maternal physiology and associated feto-placental growth. Indeed, *B. breve* induced changes in placental morphogenesis and the abundance of placental glucose and lipid transporters, which were associated with improvements in growth and metabolism of the fetus.

## Materials and Methods

### *Bifidobacterium breve* UCC2003/pCheMC

*B. breve* UCC2003/pCheMC was generated by introducing the plasmid pCheMC to electrocompetent *B. breve* UCC2003 as described previously to allow antibiotic tagging of *B. breve* for subsequent culture studies [17]. In brief, *B. breve* UCC2003 was grown until mid-log phase, chilled on ice and washed twice with ice cold sucrose citrate buffer (1mM citrate, 0.5M sucrose, pH5.8) and then electroporation of cells was carried out under the following conditions; 25MF, 200Ohms, 2KV. Transformed cells were incubated for 2 hours in Reinforced Clostridial Medium (RCM) at 37°C in a controlled anaerobic chamber then plated [18] on RCM agar plates with selective antibiotics. Colonies were sub-cultured 3 times on RCM agar plates with selective antibiotics. Antibiotics were used at the following final concentrations erythromycin 2μg/mL.

### Lyophilised *B breve*

*B. breve* was grown in De Man, Rogosa and Sharpe agar (MRS) under anaerobic conditions overnight. The bacterial cell pellet was resuspended in 10% milk powder and lyophilised in 200ml quantities. Lyophilised *B. breve* was reconstituted with 500μl PBS. Concentration of *B. breve* was 10^10^CFU/ml. All batches were tested for contamination upon reconstitution on Luria-Bertani (LB) and Brain-Heart Infusion (BHI) plates under anaerobic and aerobic conditions at 37°C. No contamination of *B. breve* was detected.

### Mice

All mouse experiments were performed under the UK Regulation of Animals (Scientific Procedures) Act of 1986. The project license PDADA1B0C under which these studies were carried out was approved by the UK Home Office and the UEA Ethical Review Committee. All mice were housed in the Disease Modelling Unit at the University of East Anglia, UK. Animals were housed in a 12:12 hour light/dark, temperature-controlled room and allowed food and water *ad libitum* (food/water intake was not recorded). Female Germ Free C57BL/6J (GF) and Specific Pathogen Free (SPF) mice aged 6-8 weeks were used for the study. GF mice were bred in germ free isolators (2 females to 1 male) and on gestational day (GD) GD9.5, pregnant mice (confirmed by weight gain) were removed from the GF isolator and transferred to individually ventilated cages. The sterility of these cages was previously tested and found to be suitable for housing GF mice for 1 week. Sterile water was changed every 2 days. We assessed responses at 2 gestational phases – the majority of studies were carried out at GD16.5, whilst the RNASeq studies utilized fetal livers harvested at GD18.5. A total of 6 SPF mice were used for GD16.5 assessments (no SPF mice were studied on GD18.5). For the GF group a total of 5 (GD16.5) and 3 (GD18.5) dams were used. For the BIF mice, a total of 6 (GD16.5) and 4 (GD18.5) dams were used.

### *B*. *breve* colonisation levels

Mice were given 100µL of reconstituted lyophilised *B. breve* UCC2003 by oral gavage (containing 10^10^ CFU/mL) at GD10, GD12 and GD14 or 100μL vehicle control (PBS, 4 % skimmed milk powder), with this dosing regimen reflecting a more realistic time frame for women who are more likely to take probiotics once their pregnancy is confirmed. At GD16.5 and GD18.5, mice were sacrificed by cervical dislocation and samples collected for molecular and histological analysis. The experimental design can be found in Figure 1A.

**Figure 1.**
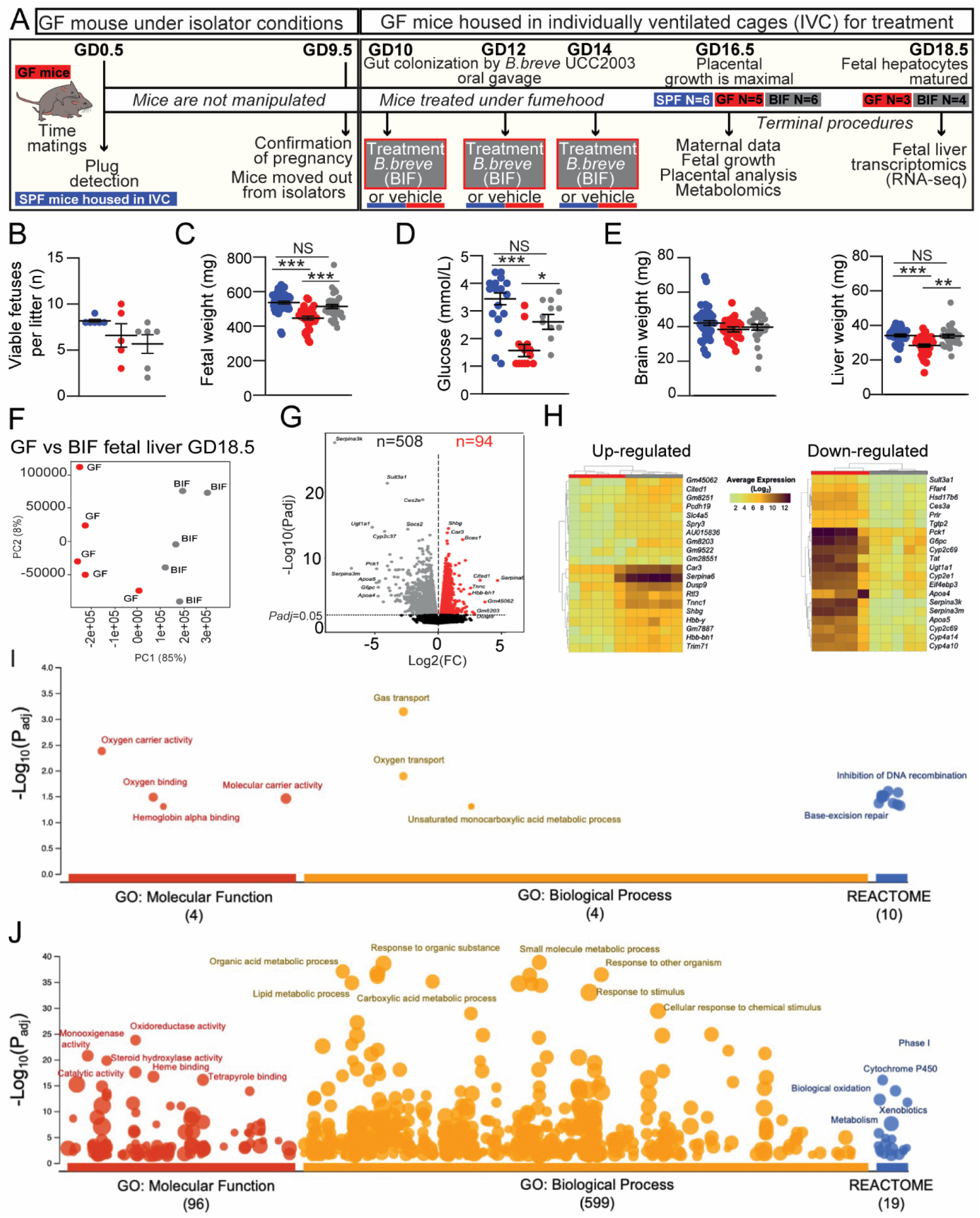
Effects of maternal gut microbiome and *B. breve* supplementation during pregnancy on fetal viability, growth and hepatic transcriptome. **(A)** Experimental design. **(B)** Number of viable fetuses per litter (One-way ANOVA with Tukey’s multiple comparison). **(C)** Fetal weight. **(D)** Circulating glucose concentrations in fetal blood. **(E)** Fetal organ weights. **(F-G)** RNA-Seq analysis of fetal liver samples obtained at GD18.5. **(F)** PCA plot and **(G)** volcano plots showing up and down-regulated differentially expressed genes (DEGs) in BIF group (compared to GF group). **(H)** Heat map of the 20 most up and down-regulated DEGs (BIF group). **(I-J)** Functional profiling (g:Profiler) on 602 DEGs. Key enriched GO terms and REACTOME pathways are shown in the figure. Fetal data were obtained on GD16.5 from: SPF (49 fetuses/6 dams), GF (33 fetuses/5 dams), BIF (34 fetuses/6 dams). Dots represent raw data for each variable assessed (individual values). However, the statistical analysis and the mean±SEM reported in the graphs were obtained with a general linear mixed model taking into account viable litter size as a covariate and taking each fetus as a repeated measure followed by Tukey multiple comparisons test (further explanations can be found in the Materials and Methods-statistical analysis section). Identification of outliers was performed with ROUT Method. RNA-seq was performed on fetal livers obtained at GD18.5 from a total of 3 GF and 4 BIF pregnant dams/litters. RNA-Seq data analysis is described in the material and methods section. (NS, not significant; *P<0.05; **P<0.01; ***P<0.001).

Faecal samples were checked for contamination and *B. breve* colonization at GD12 and GD14 and GD16. Briefly, faecal samples from GF and BIF mice were diluted in 500 µL of PBS and agitated for 30 mins at 4°C on an Eppendorf MixMate 5353 Digital Mixer Plate Shaker. The faecal solution was passed through a 0.45µm syringe filter. Faecal solution was diluted 1 in 100 and 20 µL was added to a De Man, Rogosa and Sharpe agar plate with erythromycin and incubated for 48 hours in an anaerobic chamber at 37°C. Colony forming units were counted using a click counter. In SPF animals housed in the same animal facility we have previously shown that *Bifidobacterium* represents ∼1% of the total gut microbiota [19].

### Blood hormones and circulating metabolites

Maternal blood was obtained by cardiac exsanguination immediately after cervical dislocation. Blood was centrifuged and serum collected and stored at -80ºC until further analysis. Blood glucose and serum concentrations of leptin, insulin, triglycerides, cholesterol, and free fatty acids were determined as previously reported [20]. Fetal blood glucose levels were measured with a handheld glucometer (One Touch Ultra; LifeScan) immediately after decapitation of the fetus (fetuses were selected at random).

### Placental histology

Placentas were cut in half and fixed in 4% paraformaldehyde overnight at 4ºC. Samples were washed 3 times with PBS for 15 minutes each and storage in 70% ethanol until embedding in wax. Embedded placentas were cut at 5μm thickness and stained with haematoxylin and eosin for gross morphology. Placental layer volume densities (labyrinth zone, junctional zone and decidua) were calculated using point counting and the Computer Assisted Stereological Toolbox (CAST v2.0) and converted to estimated volumes by multiplying by the weight of the placenta. For analysis of labyrinth components, sections were stained with lectin for identification of fetal endothelial vessels and with cytokeratin for trophoblasts. Further details of the double-labelling immunohistochemistry can be found elsewhere [21]. Structural analysis of the labyrinth was performed as previously described [22–24]. Briefly, fetal capillaries, maternal blood spaces and trophoblast volume densities were calculated with a point counting system in 16 random fields and their densities were then multiplied by the estimated volume of the labyrinth zone to obtain the estimated component volume. To estimate the surface density of the maternal-facing and fetal-facing interhaemal membranes, we recorded the number of intersection points along cycloid arcs in a total 20 random fields of view. Both interhaemal membrane surfaces were converted to absolute surface areas and the total surface area for exchange calculated by averaging the two absolute surface areas. Fetal capillary length densities were obtained using counting frames with two contiguous forbidden lines [24] and then converted to absolute capillary length by multiplying by the volume of labyrinth zone. Feta; capillary diameter was estimated using the equation; d=2(mean area/π)^1/2^. The interhaemal membrane barrier thickness was determined using orthogonal intercepts and measuring the shortest distance between fetal capillaries and the closest maternal blood spaces at random starting locations (at least 99) within the labyrinth zone [24].

For the analysis of placental glycogen, sections were stained with Periodic acid–Schiff (Sigma-Aldrich) previous incubation with 0.5% periodic acid (Thermo Fisher Scientific). Sections were counterstained with Fast-green (Sigma-Aldrich) and digitalized with the nanozoomer scanner (Hamamatsu). Analysis of placental glycogen accumulation was performed with Image J and conducted blinded to experimental groups. TUNEL staining for placental cell death was performed using the TUNEL Assay Kit - HRP-DAB (Abcam, ab206386) following manufacturer instructions except for the counterstaining which was substituted for Nuclear Fast Red (Vector). Sections were digitalized using a nanozoomer scanner (Hamamatsu) and the amount of apoptosis in the labyrinth zone was calculated in 5 random areas (x20 magnification) and analysed by Image J software.

### Western blotting

Protein extraction was performed with RIPA buffer as described previously [25]. Lysates were separated by SDS-PAGE and incubated with antibodies against p-MAPK (Thr202/Tyr204) (Cell Signalling, 4370; 1/1000), t-MAPK 44/42 (Cell Signalling, 4695; 1/1000), DLK-1 antibody (Abcam, ab21682; 1/1000), p-P38MAPK (Cell Signalling, 4511; 1/1000) and t-P38MAPK (Cell Signalling, 8690; 1/1000). Reactive bands were detected by chemiluminescence (Thermo Scientific, Scientific SuperSignal West Femto) and quantified by Image J software. Proteins were normalized to Ponceau S Staining [26].

### RNA extraction and qPCR

Extraction of RNA from micro-dissected placental labyrinth zones was performed with RNeasy Plus Mini Kit (Qiagen) and reverse transcribed using the High Capacity cDNA RT Kit minus RT inhibitor (Applied Biosystems) according to manufacturer’s instructions. Samples were analysed using MESA Blue SYBR (Eurogentec) and primers (See Table S1) were synthesized by Sigma-Aldrich. The expression of each gene was normalized to the geometric mean expression of two reference genes *Hprt* and *Ubc*, which remained stably expressed across the groups. Analysis was performed using the 2-ΔΔCt method [27].

### Sequence pre-processing, Differential Gene Expression (DGE) analysis and Functional enrichment analysis

Fetal liver RNA on GD18.5 was extracted using the RNeasy Plus Mini Kit (Qiagen). Purified RNA was quantified, and quality controlled using RNA 6000 Nano kit on a 2100 Bioanalyser (Agilent). Only samples with RIN values above 8 were sequenced. RNA sequencing was performed at the Wellcome Trust Sanger Institute (Hinxton, UK) on paired-end 75 bp inserts on an Illumina HiSeq 2000 platform. Isolated RNA was processed by poly-A selection and/or Ribo-depletion. RNA sequence pre-processing and DGE analysis was performed as previously described with slight modifications [28]. Briefly, FASTQ reads were initially quality-filtered using fastp v0.20.0 with options -q 10 (sequence reads with phred quality <10 were discarded). Subsequently, sequence reads for each sample were merged (merge-paired-reads.sh) and followed by rRNA sequence filtering via SortMeRNA v2.1 based on SILVA rRNA database optimised for SortMeRNA software [29, 30]. Filtered reads were then unmerged (unmerge- paired-reads.sh) and ready for transcript quantification. Transcript mapping and quantification were then performed using Kallisto v0.44.0 [31]. *Mus musculus* (C57BL/6 mouse) cDNA sequences (GRCm38.release-98_k31) retrieved from Ensembl database were indexed with Kallisto utility *index* at default parameter and was used for following transcript mapping and abundance quantification via Kallisto utility *quant* at 100 bootstrap replicates (-b 100) [32].

RNA raw counts were subjected (Kallisto outputs) to DGE analysis, which was performed using R library *Sleuth* (v0.30.0) [33]. Transcripts were then mapped to individual genes using Ensembl BioMart database (GRCm38.p6) with function *sleuth_prep* and option *gene_mode = TRUE*. Genes with an absolute log_2_ (fold change) >1.0 and q value <0.05 (p-adjusted value; based on Wald test statistics) were considered to be differentially regulated [34]. DGE statistics were plotted via functions within package *Sleuth*. Finally, functional enrichment analysis was performed using g:Profiler webtool g:GOst based on organism *Mus Musculus* species [35]. Briefly, a list of DGEs (Ensembl IDs) was uploaded to g:GOst, then selected ‘GO molecular function’, ‘GO biological process’ and ‘Reactome’ in the ‘data sources’. Significance threshold was set at 0.001 (g:SCS threshold).

### Metabolite extraction, Nuclear Magnetic Resonance (NMR) spectroscopy and metabolite quantification

Extraction of metabolites from fetal liver, placenta and maternal caecum contents were performed as previously described as a standard protocol [36]. For caecal samples, frozen materials (stored at -80°C prior to analysis) were weighed ∼50mg before the addition of 600μL of faecal water phosphate buffer solution. The faecal water phosphate buffer was prepared as follows: add 0.51g NaH_2_PO_4_.H_2_O and 2.82g K_2_HPO_4_ to 200 mL D_2_O (Deuterium Oxide; Merck). To this, 34.5mg TSP (Trimethylsilyl propanoic acid; used as NMR standard) and 100mg NaN_3_ (Merck) were added [37]. Next, the mixture was centrifuged for 10 min at 17,000 x g before transferring the mixture to an NMR tube (Merck) for subsequent NMR analysis.

For liver and placenta samples (stored at -80°C prior to analysis), frozen fresh tissue (∼20-45 mg) was placed into a 2ml sterile microcentrifuge tube pre-loaded with ∼15-20 glass beads (Merck) while 200μL of ice-cold methanol (Fisher Scientific) and 42.5μL of ultra-pure cold water were added to it and vortexed. Tissue was disrupted via a tissue lyser (Qiagen) for 2 × 2 mins. 100μL of ice-cold chloroform (Merck) was then added and vortexed. 100μL of ice-cold chloroform and 100μL of ultra-pure cold water were added to the mixture, and kept on ice for 15 minutes. Liquid was then transferred into a new sterile microcentrifuge tube and centrifuged for 3 minutes at 17,000 × g. The top aqueous phase was transferred into a new microcentrifuge tube and speed-vacuumed for 30 minutes at 50°C and 30 minutes without heating prior to reconstitution with faecal water phosphate buffer solution at 600 μL. The mixture was then moved to an NMR tube (Merck) for subsequent NMR analysis. Metabolites from culture media Brain Heart Infusion (BHI; Oxoid) and spent media (BHI cultured with *B. breve* UCC2003 for 48 h) were extracted as follows: 400μL of medium was transferred into a sterile microcentrifuge tube with the addition of 200μL faecal phosphate buffer and mixed well. The mixture was then moved to an NMR tube (Merck) for further NMR analysis.

Samples in NMR tubes were subsequently subjected to NMR spectroscopy. The ^1^H NMR spectra were recorded at 600 MHz on a Bruker AVANCE spectrometer (Bruker BioSpin GmbH, Germany) running Topspin 2.0 software. The metabolites were then quantified using the software Chenomx® NMR Suite 7.0™.

### Statistical analysis

All statistical analysis and sample size are shown in each figure/table and in the corresponding figure/table legends. Only samples from viable fetuses were analysed. No statistical analysis was used to pre-determine sample size and samples were assigned code numbers and, were possible, analysis was performed in a blinded fashion. Statistical calculations were performed using the GraphPad Prism software (GraphPad v9, San Diego, CA), SAS/STAT 9.0 (Statistical System Institute Inc. Cary, NC, USA) and RStudio Version 1.4.1106 (RStudio Boston, MA) with R Version 4.0.3 (Vienna, Austria). Data reported as mean±SEM. Morphometric parameters of mother, litter size and western blot data were analysed by one-way ANOVA followed by Tukey post hoc test. Feto-placental weights, placental stereological measurements and placental Lz gene expression levels were analysed with a general linear mixed model, taking into account viable litter size as a covariate and taking each fetus as a repeated measure of the mother. In this statistical analysis, fetuses and placentas per litter are nested within litters[38]. Identification of outliers was performed with ROUT Method. For metabolomics, differences between individual metabolites between the three groups were tested with a Kruskal-Wallis test using the kruskal.test function with correction for multiple comparisons applied using the Benjamini & Hochberg false discovery rate method using the p.adjust function. Pairwise comparisons between the three groups were carried out with a Dunn’s test on individual metabolites significantly different after correction for multiple comparisons using the dunnTest function in the FSA package. The level of significance for all statistical tests used in this study was set at *P*<0.05. All figures in the manuscript show individual values (raw data). However, P values and mean±SEM within the graphs analysed by the general linear mixed model were corrected for repeated measures. Graphs containing the individual dots and graphs with corrected mean±SEM were generated with Graphpad and merged with Adobe Illustrator.

## Results

### Germ-free mice treated with B. breve have altered body composition and caecum metabolic profile

To assess whether maternal microbiota can influence feto-placental growth, GF mice were treated orally with *B. breve* UCC2003 from day 10 of gestation (treatment on days 10, 12 and 14; i.e. BIF group), and compared to GF and SPF dams (for experimental overview see Figure 1A). Timing and dosing were based on the fact that levels of *Bifidobacterium* rise throughout pregnancy (5) (colonization levels during pregnancy can be found in Figure S1). Previous work has indicated three consecutive doses of *B. breve* UCC2003 facilitates stable gut colonization, with the advantage of also avoiding repeated handling of the mice, which may induce spontaneous abortions [28, 39]. In addition, from a translational point of view, we also wanted to correlate our animal model with potential future supplementation studies in women at the point pregnancy is confirmed.

Maternal body composition differed between groups with GF and BIF mice showing increased digestive tract weight and lower pancreas mass compared to SPF mice. GF and BIF mice had similar circulating concentrations of glucose and insulin to SPF mice (Table 1). Compared to SPF mice, treatment with *B. breve* reduced maternal gonadal fat depot, liver, and spleen weights in BIF mice. No differences were observed in the circulating concentrations of leptin, cholesterol, triglycerides, or free fatty acids in maternal serum (Table 1).

**Table 1.** Effects of maternal gut microbiome and *B. breve* administration during pregnancy on maternal body composition, circulating metabolites and hormones in maternal serum, and metabolites in caecum on day 16.5 of gestation. Body composition and metabolites/hormones in serum were analyzed by one-way ANOVA followed by Tukey multiple comparisons test. Metabolites in maternal caecum were analysed by Kruskal-Wallis test followed by multiple comparisons using the Benjamini & Hochberg false discovery rate method and Dunn’s test. ROUT test was used for identification of outlier values. The level of significance was set at *P*<0.05. NS: not significant. Data presented as mean±SEM. The number of dams used for each group is annotated on the table and only data from dams at day 16.5 of gestation were used.

|  | SPF (n=6) | GF (n=5) | BIF (n=6) | SPF vs GF | SPF vs BIF | GF vs BIF |
| --- | --- | --- | --- | --- | --- | --- |
| Hysterectomy weight (g) | 26.01±0.91 | 27.87±0.78 | 27.17±0.80 | NS | NS | NS |
| Digestive Tract (g) | 2.76±0.03 | 6.83±0.32 | 7.25±0.63 | <0.0001 | <0.0001 | NS |
| Caecum (g) | 0.66±0.03 | 3.47±0.25 | 3.96±0.41 | <0.0001 | <0.0001 | NS |
| Small intestine (g) | 1.66±0.03 | 2.65±0.11 | 2.59±0.14 | <0.0001 | <0.0001 | NS |
| Pancreas (mg) | 315.40±30.12 | 183.40±24.74 | 190.60±38.71 | 0.041 | 0.044 | NS |
| Gonadal fat (mg) | 433.10±43.20 | 297.0±37.02 | 272.0±27.35 | NS | 0.016 | NS |
| Liver (g) | 2.09±0.10 | 1.79±0.05 | 1.55±0.08 | NS | 0.001 | NS |
| Spleen (mg) | 117.90±2.80 | 91.76±10.60 | 83.03±6.72 | NS | 0.0127 | NS |
| Glucose (mmol/L) | 8.08±0.78 | 8.38±1.18 | 8.88±0.74 | NS | NS | NS |
| Insulin (µg/L) | 0.12±0.004 | 0.19±0.05 | 0.20±0.06 | NS | NS | NS |
| Leptin (pg/mL) | 2465±177.1 | 2739±486 | 2425±303 | NS | NS | NS |
| Cholesterol (mmol/L) | 1.33±0.03 | 1.56±0.08 | 1.41±0.09 | NS | NS | NS |
| Triglycerides (mmol/L) | 1.54±0.08 | 1.79±0.14 | 1.50±0.11 | NS | NS | NS |
| Free Fatty Acids (µmol/L) | 890.6±101.3 | 1440±362 | 1092±114.5 | NS | NS | NS |
| **Maternal caecum metabolites** |  |  |  |  |  |  |
|  | SPF (n=3) | GF (n=4) | BIF (n=5) | SPF vs GF | SPF vs BIF | GF vs BIF |
| Butyrate (mmol/Kg) | 12.47±7.97 | 0.00±0.00 | 0.00±0.00 | 0.006 | 0.008 | NS |
| Citrulline (mmol/Kg) | 0.30±0.06 | 0.00±0.00 | 0.00±0.00 | 0.006 | 0.008 | NS |
| Fucose (mmol/Kg) | 0.08±0.01 | 0.00±0.00 | 0.00±0.00 | 0.006 | 0.008 | NS |
| Isobutyrate (mmol/Kg) | 0.41±0.24 | 0.00±0.00 | 0.00±0.00 | 0.006 | 0.008 | NS |
| Isovalerate (mmol/Kg) | 0.09±0.01 | 0.00±0.00 | 0.00±0.00 | 0.006 | 0.008 | NS |
| Malonate (mmol/Kg) | 0.09±0.02 | 0.00±0.00 | 0.00±0.00 | 0.006 | 0.008 | NS |
| Methylamine (mmol/Kg) | 0.05±0.02 | 0.00±0.00 | 0.00±0.00 | 0.006 | 0.008 | NS |
| Propionate (mmol/Kg) | 4.48±1.99 | 0.00±0.00 | 0.00±0.00 | 0.006 | 0.008 | NS |
| Trimethylamine (mmol/Kg) | 0.06±0.007 | 0.00±0.00 | 0.00±0.00 | 0.006 | 0.008 | NS |
| Valerate (mmol/Kg) | 0.57±0.27 | 0.00±0.00 | 0.00±0.00 | 0.006 | 0.008 | NS |
| 2.methylbutyrate (mmol/Kg) | 0.05±0.01 | 0.00±0.00 | 0.00±0.00 | 0.006 | 0.008 | NS |
| 5.Aminopentanoate (mmol/Kg) | 0.33±0.14 | 0.00±0.00 | 0.00±0.00 | 0.006 | 0.008 | NS |
| Acetate (mmol/Kg) | 35.49±20.66 | 0.55±0.06 | 3.22±1.39 | 0.006 | 0.129 | 0.094 |

Metabolomics analysis in maternal caecum samples indicated that the concentration of 13 out of 81 metabolites were significantly altered (Table 1 and Table S2). Acetate was influenced by *B. breve* (Table 1), with BIF dams having intermediate concentrations compared to SPF and GF mice (the low levels of acetate detectable in GF mice, most likely originated from the diet and/or are host-derived). These findings suggest that acetate producing *B. breve* and the wider gut microbiota may exert selective effects on maternal metabolic (gonadal fat depot and liver) and immune organs (spleen).

### Maternal gut microbiota and B. breve regulate fetal growth by controlling fetal glycaemia and hepatic transcriptome

The three experimental groups had similar numbers of viable fetuses per litter, although GF and BIF groups showed a higher variability compared to the SPF group (Figure 1B). Compared to SPF and BIF mice, GF fetuses were growth restricted, hypoglycaemic and had reduced liver weight, but had preserved brain size (Figure 1C-E). As the liver is a key organ for glucose storage and production and fetuses from BIF mice had heavier livers and improved glycaemia, we next determined if there were changes in the fetal hepatic transcriptome (livers were collected from a small cohort of mice on GD18.5, when fetal liver function is particularly active prior to term. Indeed, mouse fetal hepatocytes are mature from GD18.5, when they present a similar gene expression pattern to those in the postnatal liver [40]). A total of 602 genes were differentially expressed, with 94 significantly up- regulated and 508 down-regulated genes in BIF group, when compared to GF group (Figure 1F-H). Functional enrichment analysis indicated that pathways involved in haemoglobin and oxygen transport-binding were significantly upregulated in the fetal livers of BIF mice (Figure 1I and Table S3). In contrast, many metabolic pathways were downregulated in response to *B. breve* administration, including carboxylic acid and lipid metabolic processes, steroid hydroxylase activity, fatty acid metabolism and response to glucocorticoid (Figure 1H; Table S3). Therefore, maternal *B. breve* appears to exert changes in fetal hepatic function with implications for fetal growth.

### Maternal gut microbiota and B. breve control placental morphogenesis

To further understand the links between the maternal gut microbiota and the regulation of fetal growth, we assessed placental structure (performed on GD16.5, when placental growth in mice is maximal [24]). When compared to SPF mice, placentas were lighter in GF and BIF mice (Figure 2A). Placental efficiency, defined as the grams of fetus produced per gram of placenta, was significantly improved in the BIF group compared to GF mice (Figure 2B). Analysis of placental compartments showed that lack of maternal gut microbiota significantly hampered growth of the placental labyrinth transport zone (Lz), without compromising the endocrine junctional zone or decidua volumes (Figure 2C). It also did not affect placental glycogen storage (Figure 2D) or the volume of the trophoblast (Figure E-F). Analysis of maternal blood spaces revealed that GF and BIF groups had reduced spaces compared to SPF mice, while the volume and the length of fetal capillaries were significantly reduced in the GF compared to SPF (Figure 2F-G). Similarly, surface area for exchange of the Lz was significantly decreased in GF compared to SPF mice (Figure 2H). The barrier between maternal and fetal blood was also determined to be thinner in BIF *versus* GF mice (Figure 2I). Lz apoptosis levels were similar between groups (Figure 2J).

**Figure 2.**
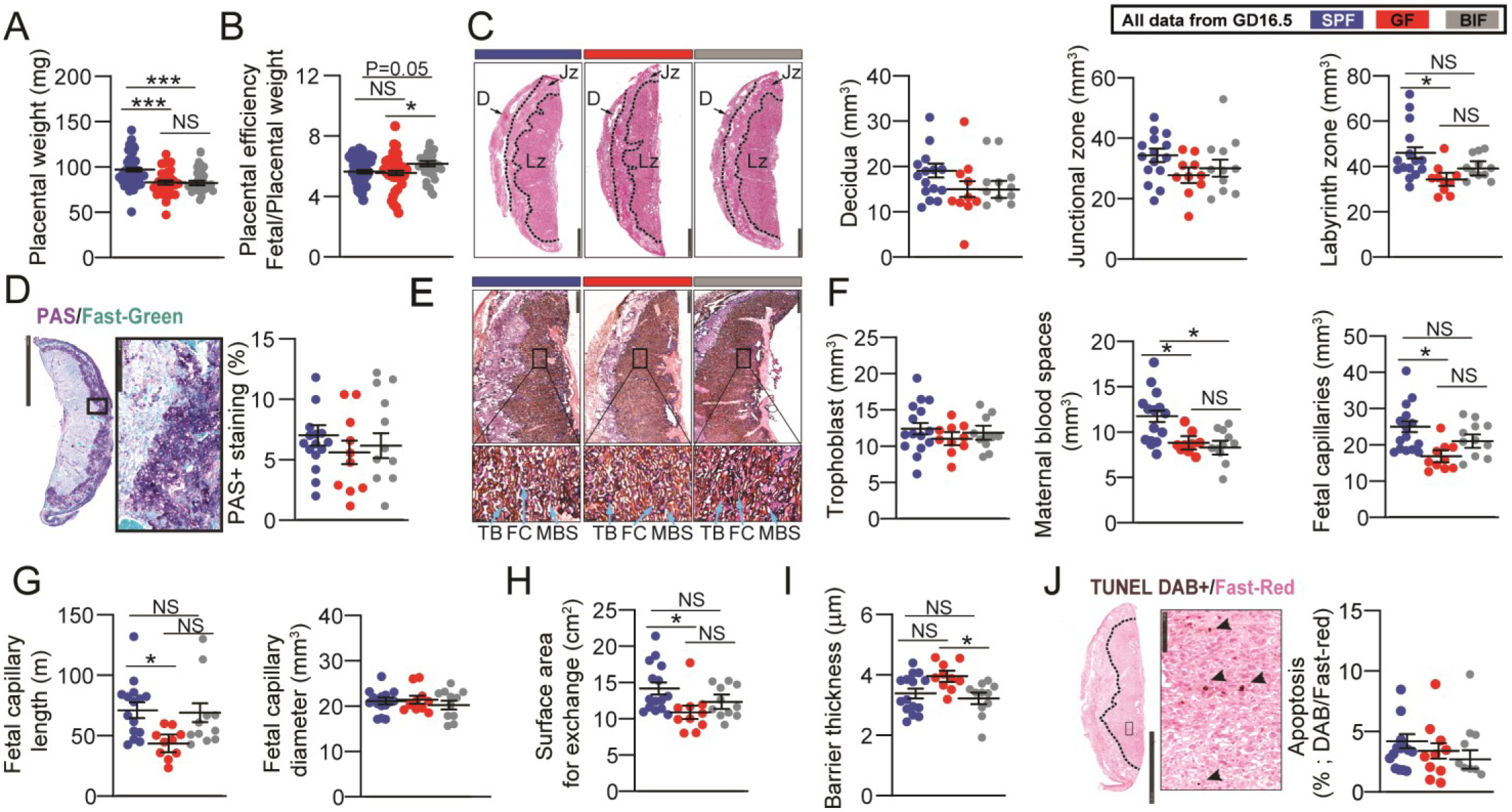
Effects of maternal gut microbiome and *B. breve* supplementation during pregnancy on placental structure on day 16.5 of gestation. **(A)** Placenta weight. **(B)** Placental efficiency determined by dividing fetal by placental mass. Scale bar= 1mm. **(C)** Placental regional analysis. **(D)** Representative staining of placental glycogen with PAS and glycogen abundance. Scale bar= 2.5mm and 250μm. **(E)** Representative image of lectin and cytokeratin staining for labyrinth zone structural quantification. Scale bar= 500μm and 50μm. **(F-I)** Stereological parameters determined in placental labyrinth zone. **(J)** Representative image of TUNEL staining for apoptosis quantification in labyrinth zone. Scale bar= 2.5mm and 100μm. All data were analyzed by general linear mixed model, taking into account litter size as a covariate and taking each fetus as a repeated measure followed by Tukey multiple comparisons test. ROUT test was used for identification of outlier values. Dots represent raw data (individual values). However, the statistical analysis and the mean±SEM reported within the graphs were obtained with the general linear mixed model (further explanations can be found in the Materials and Methods statistical analysis section). Placental weight-efficiency was obtained from: SPF (49 fetuses/6 dams), GF (33 fetuses/5 dams), BIF (34 fetuses/6 dams). Laboratorial analysis was performed with: SPF (14-15 placentas from 6 dams), GF (10 placentas from 5 dams) and BIF (9-11 placentas from 6 dams). Only placentas collected on day 16.5 of gestation were analysed. One to three placentas per litter were randomly selected and used for assessment. Placentas were analysed blind to the experimental groups. (NS, not significant; *P<0.05; ***P<0.001). Abbreviations: D (decidua), Jz (junctional zone), Lz (labyrinth zone), TB (trophoblasts), FC (fetal capillaries), MBS (maternal blood spaces).

To define the molecular mechanisms behind the changes in the Lz, we quantified the expression of select genes in micro-dissected Lz. The angiogenic factor *Vegf* was similarly expressed between groups (Figure 3A). However, the expression of signalling pathways involved in cell proliferation and growth, namely the MAPK pathway, was significantly altered by changes in maternal gut microbiota; *Mapk1* was shown to be increased in both GF and BIF, while *Mapk14* (also known as *p38Mapk*) was revealed to be specifically up-regulated in the Lz of BIF mice. In addition, *Dlk1* and *Igf2P0*, which are key genes implicated in metabolism and Lz formation, were significantly up-regulated in the BIF group compared to GF mice. The expression of *Akt* did not vary with group (Figure 3A). As informed by western blotting, activation of ERK was reduced in the placental Lz of GF compared to SPF mice, and this effect was reversed by BIF (Figure 3B). p38MAPK protein activity was similar between groups. DLK1 protein level was also lower in GF compared to SPF mice. However, BIF increased DLK1 protein levels when compared to both SPF and GF mice (Figure 3B). Overall, these findings suggest that the maternal gut microbiota, and *B. breve*, regulate the development of the mouse placental Lz via modulation of specific cell growth and metabolic genes/pathways.

**Figure 3.**
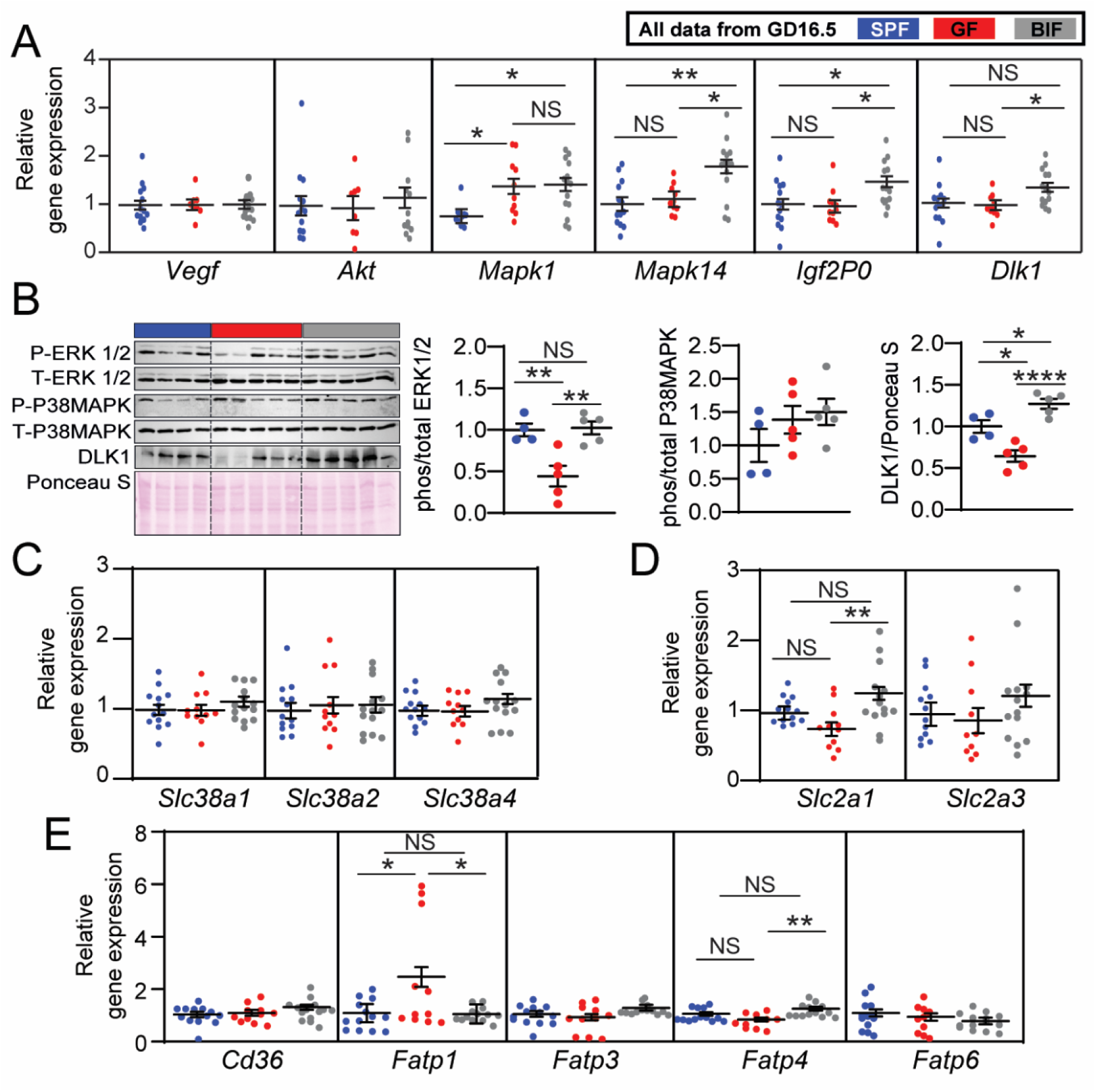
Effects of maternal gut microbiome and *B. breve* supplementation during pregnancy on placental gene and protein levels on day 16.5 of gestation. **(A)** Gene expression levels in micro- dissected labyrinth zones. **(B)** Immunoblots and protein quantification by Western blot in micro- dissected labyrinth zones. **(C-E)** Gene expression levels in micro-dissected labyrinth zones for amino- acids, glucose and lipid transporters. Western blot data was analysed by one-way ANOVA. qPCR data were analyzed by general linear mixed model, taking into account litter size as a covariate and taking each fetus as a repeated measure followed by Tukey multiple comparisons test. ROUT test was used for identification of outlier values. Dots represent raw data (individual values). However, the statistical analysis and the mean±SEM reported within the graphs (for qPCR data) were obtained with the general linear mixed model (further explanations can be found in the Materials and Methods statistical analysis section). Gene expression analysis was performed with: SPF (13 placentas from 6 dams), GF (11 placentas from 5 dams) and BIF (14 placentas from 6 dams). Protein quantification was performed with: SPF (4 placentas from 4 dams), GF (5 placentas from 5 dams) and BIF (5 placentas from 5 dams). Only placentas collected on day 16.5 of gestation were analysed. For qPCR, one to three placentas per litter were assessed and selection of the samples was conducted at random. For protein expression analysis, 1 placenta per litter was selected. (NS, not significant; *P<0.05; **P<0.01).

### Maternal gut microbiota and B. breve controls key placental nutrient transporters

To better understand the changes in fetal growth and glycemia between groups, we quantified the expression of selected amino acid, glucose and lipid transporters in the Lz. We found no difference in the expression of system A amino acid transporters (*Slc38a1, Slc38a2, Slc38a4*) between groups (Figure 3C). However, the key glucose transporter *Slc2a1* was up-regulated in the Lz of BIF mice compared to GF mice (*Slc2a3* mRNA levels were similar between groups; Figure 3C). Fatty acid transporters were also altered, with increased levels of *Fatp1* in the GF group compared to SPF and BIF, while *Fatp4* was increased in the BIF group compared to the GF (Figure 3D; *Cd36* and *Fatp3,6* expression levels were unaltered). Collectively, these data suggest that maternal gut microbiota, and *B. breve*, could regulate fetal growth by inducing changes in the expression of key nutrient transporters within the placenta.

### Differences in placental labyrinth growth are linked to an altered placental metabolome

To gain further mechanistic understanding of the changes observed in the placental Lz and fetal liver, we analysed >80 metabolites at GD16.5 (Figure 4 and Table S2). We found 5 metabolites significantly altered in the placental Lz (Figure 4). 2-Aminoadipate in the Lz was very low in GF/BIF groups as well as in fetal livers (Figure 4A). Treatment with *B. breve* significantly reduced the concentrations of acetylcarnitine and carnitine in Lz tissue compared to SPF placentas, but not in fetal livers (Figure 4B- C). Levels of formate in placental Lz were significantly elevated in both GF and BIF compared to SPF mice (Figure 4D), with a similar trend (although not significant) in fetal liver samples. Acetate was also altered in the Lz (Figure 4E), with concentrations significantly lower in the SPF compared to the GF group, whilst BIF samples showed intermediate levels (although these levels were much lower than observed in the maternal caecum). Similar to formate, concentrations of acetate in fetal liver followed similar directions to the Lz, yet were not statistically different between groups. These data suggest that maternal gut microbiota, and *B. breve*, regulate the fetal and placental growth via modulation of the placental Lz metabolome.

**Figure 4.**
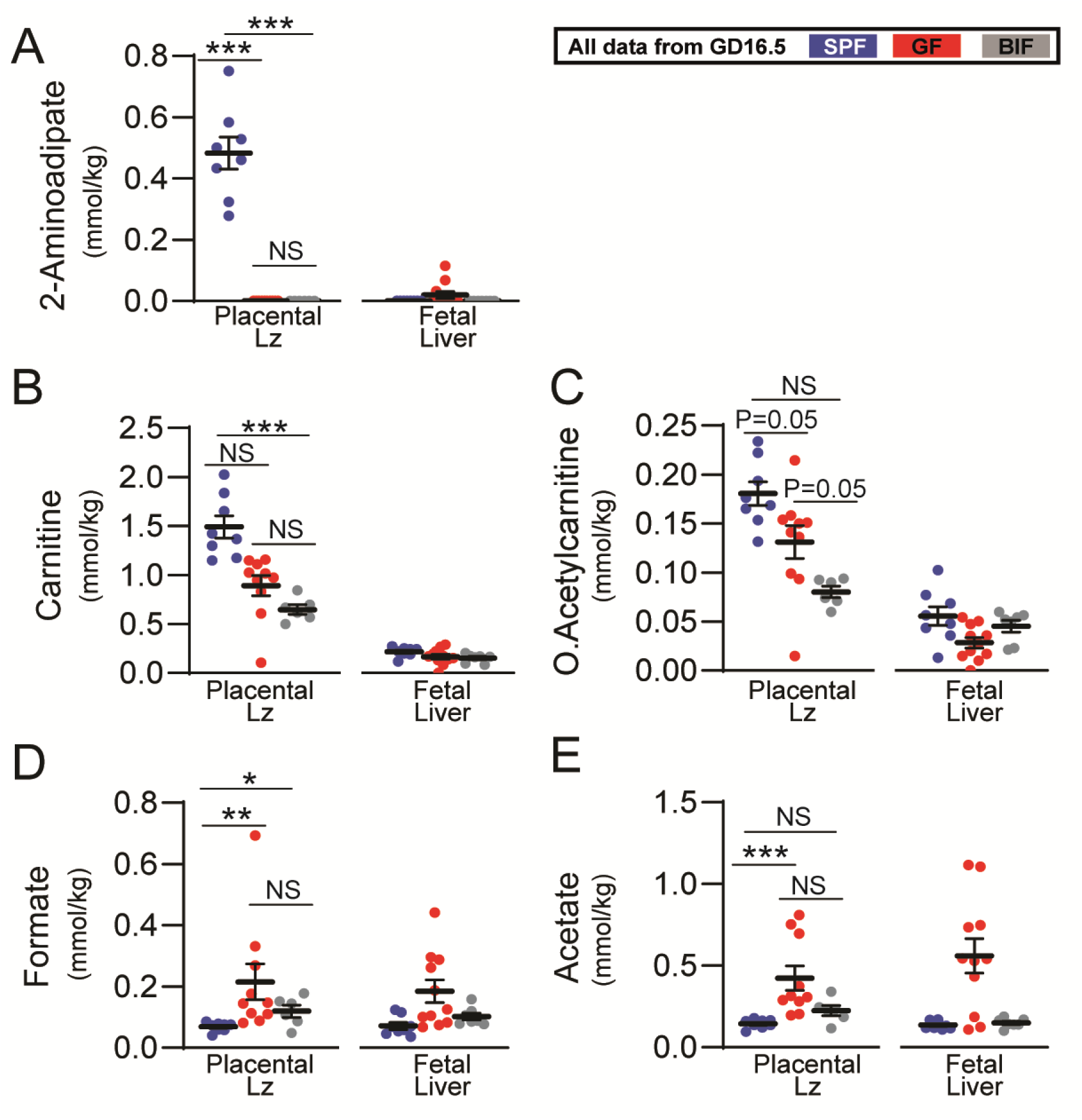
Metabolomic profiling of placental labyrinth zone and fetal liver on day 16.5 of gestation. Data were analysed by Kruskal-Wallis test followed by multiple comparisons using the Benjamini & Hochberg false discovery rate method and Dunn’s test. ROUT test was used for identification of outlier values. Different superscripts indicate significant differences between groups (a *versus* b, P<0.05). Data presented as mean±SEM. Number of litters analysed per group: SPF (8 placentas-livers from 4 dams), GF (8-7 placentas-livers from 4-5 dams), BIF (6-7 placentas-livers from 5 dams). Only tissues collected at GD16.5 were analysed. Selection of the samples was conducted at random. (NS, not significant; *P<0.05; **P<0.01; ***P<0.001).

## Discussion

In this study, we demonstrate that the maternal gut microbiota and the microbiota member *B. breve* regulate feto-placental growth. To the best of our knowledge, this is the first demonstration of a maternal gut bacterium remotely controlling placental structure and nutrient transporters, with important implications for fetal glycaemia and fetal growth. We observed that the effects of *Bifidobacterium* are partially mediated by altered metabolites in maternal caecum and in placental Lz tissue, with alterations in the expression of key genes in the placental Lz and fetal liver.

*Bifidobacterium* is the dominant microbiota member in vaginally delivered, breast-fed infants, with certain species and strains known to stimulate and aid the maturation of the immune system [41]. *B. breve* UCC2003 also regulates responses at the gut barrier, inducing homeostatic epithelial cell programming, and protecting against inflammatory insults [28, 42]. Importantly, pregnancy is accompanied by increasing *Bifidobacterium* abundance in the gut of women and mice [5] and alterations in the abundance of *Bifidobacterium* are linked to the development of serious pregnancy complications like preeclampsia [43]. Recently, it has been demonstrated that the maternal gut microbiota regulate embryonic organ growth by promoting fetal neurodevelopment [44]. Our study shows that maternal gut microbiota induces changes in fetal organogenesis and that *B. breve* supplementation restored fetal glycaemia and liver weight. In this regard, fetal brain weight was unaltered in the three experimental groups, whilst liver mass was drastically reduced only in the GF group. Together, these results suggest that untreated GF fetuses prioritize the growth of the brain at expense of the liver. This fetal strategy, known as the ‘brain sparing effect’, is a protective mechanism to preserve oxygenation and nutrient delivery to the brain in situations of placental insufficiency [45– 48]. Our RNAseq analysis shows an upregulation of genes involved in oxygen transport and haemoglobin binding, and downregulation of metabolic pathways like steroid hydroxylase activity, carboxylic acid binding or fatty acid metabolism in the BIF group. These data therefore suggest that fetal defenses against growth retardation were better in the BIF group compared to the GF group, and that the downregulation of the metabolic pathways could be due to the fact that BIF fetuses already achieved their hepatocyte maturation or their maximum liver growth potential earlier than the GF group. In fact, *B. breve* supplementation restored fetal glycaemia and weight, achieving similar values to that seen for SPF fetuses.

Previous *in vivo* studies show different strains of *Bifidobacterium* (including *B. breve*) modulate glucose handling [49], with this genus consistently associated with potential protection against human metabolic disorders e.g. type 2 diabetes [50, 51]. Our observations of reduced maternal gonadal fat mass and maternal liver weight in *B. breve*-treated dams compared to SPF dams, suggest that *Bifidobacterium*, or *B. breve* metabolites, could affect responses of key organs in the mother, and subsequently impact fetal resource allocation. *B. breve* UCC2003 appeared to induce changes in the metabolite milieu, including carnitine and acetate in the maternal caecum and/or placenta, which could be determinant for the effects observed on fetal growth. Carnitine is well-known for mediating the transport of fatty acids into mitochondrial matrix for fatty acid β-oxidation and BIF placentas had lower concentrations of acetylcarnitine and carnitine compared to SPF. These results suggest a potential greater reliance on these compounds for energy production, or enhanced transfer of these fatty acids to the fetus [52]. On the other hand, acetate is a major bifidobacterial fermentation by- product, which directly mediates glucose homeostasis through the free fatty acid receptor 2 [53] and epithelial cell responses. Previous work in adult mice suggests that elevation of gut acetate levels due to *Bifidobacterium* treatment plays a key role in regulating glucose handling systemically and reduces visceral fat accumulation [54]. Acetate also exerts systemic metabolic [55, 56] and immunological effects [57]. More generally, microbial-derived short-chain fatty acids (SCFAs) modulate multiple host physiological systems and during pregnancy are associated with maternal gestational weight, neonatal length and body weight, and protection against allergic airway disease in the developing fetus [58, 59]. Acetate crosses the placenta [59], so in our model, the elevated maternal *B. breve*-derived acetate may exert effects on feto-placental growth in three potential ways. Firstly, higher maternal caecum acetate concentrations in SPF and *B. breve* supplemented dams vs. GF dams could indicate maternal effects, through interactions within the maternal gut mucosa and subsequent impact on maternal organs (liver, adipose and spleen). Secondly, effects on the placenta, through the potential use of acetate for cellular metabolism, growth and function. Finally, effects on fetal metabolism following transport of acetate across the placenta to the fetus. Compared to the maternal caecum, levels of acetate were relatively low in the placental Lz and fetal liver (for all 3 groups). This suggests that *B. breve* (and SPF microbiota- derived acetate) may be used to support anabolic processes *in utero* (hence the very low levels detected). Moreover, the observed modulation of immune-associated pathways in the fetal liver, including those associated with G protein-coupled receptor signalling (e.g. *Dusp9*), also indicates a role for direct acetate-associated responses [60]. Further work is required to fully understand the mechanisms behind the differences observed in maternal organs between the SPF and BIF groups (liver, adipose and spleen) and how these changes impact on materno-fetal resource allocation. Thus, future work should assess the ontogeny of these changes and incorporate an additional pregnant SPF group treated with *B. breve* to fully understand the chemical, endocrine and metabolic interactions occurring between *B. breve*, maternal organs (gut, liver, adipose and spleen) and fetal metabolism.

Administration of *B. breve* significantly reduced the interhaemal membrane barrier thickness of the placenta (compared to GF group), which may facilitate exchange of nutrients and gases. Previous work has shown that the barrier thickness is regulated by *Igf2P0*, as *Igf2P0*/knockout mice have increased thickness of the exchange barrier and reduced passive permeability of the placentas [61]. In our case, *Igf2P0* was significantly elevated in the BIF group compared to the SPF and GF groups and although *in vivo* functional assays evaluating the passive and active transport of solutes are required to verify the implications of this effect on fetal nutrient allocation (e.g. performing unidirectional maternal–fetal transfer assays using ^51^Cr-EDTA or glucose and amino acid analogues, ^3^H-MeG or ^14^C-MeAIB [23, 61, 62]), this result explains, in part, the improvement in fetal weight observed in the BIF group. Moreover, IGFs have been implicated in the regulation of glucose transporters in a variety of organs by utilizing signalling pathways like PI3K/AKT and MAPK [63], and among the different nutrient transporters evaluated, *Slc2a1* mRNA levels were significantly elevated in the BIF group compared to GF group. The other two transporters that were altered, *Fatp1* and *Fatp4*, changed in opposite directions suggesting that *B. breve* could modulate the expression of these two transporters in different ways depending on the direction and magnitude of fatty acid flux at the placental Lz [64, 65]. The divergent expression of *Fatp1* and *Fatp4* in the BIF compared to GF group may also be linked to intracellular carnitine utilization, as *Fatp1* can interact with carnitine palmitoyltransferase 1 to promote fatty acid transport into mitochondria [66].

Maternal gut microbiota affected placental structure and its vascular bed, which is required for adequate fetal growth [23]. The mechanisms governing these structural changes could be partially mediated by changes in the expression *Igf2P0* and *Dlk1* (two important imprinted genes in placental physiology [67]), and via MAPK upregulation. In addition to the changes above described for barrier thickness, deletion of *Igf2P0* results in feto-placental growth restriction in association with reduced placental surface area for exchange and fetal capillary volume (reviewed by [68]); parameters that were significantly affected in the SPF vs GF groups, and partially restored by *B breve* administration. *Dlk1*, a non-canonical ligand of the Notch signalling pathway localized to the endothelial cells of fetal capillaries in the placental Lz, regulates placental vascularisation and branching morphogenesis [69] and both, IGF2 and DLK1, can mediate cellular actions via the MAPK pathway [70–72]. We observed no differences in the mRNA levels of *Dlk1* between SPF and GF mice. However, at the protein level, DLK1 in the Lz was controlled by the maternal gut microbiota and more specifically by *B. breve*. Similarly, *Mapk1* levels were increased in both GF and BIF groups compared to the SPF, but at the protein level, ERK activity was lower in the GF but not BIF group when compared to the SPF group, suggesting once again that *B. breve* is involved in the regulation of DLK-MAPK signaling. Another important signaling pathway for embryogenesis and placental Lz angiogenesis and vascular remodeling is p38MAPK (encoded by the gene *Mapk14*) [73]. This pathway has been linked to environmental stresses and inflammatory cytokines [74]. However, p38MAPK also regulates many normal cellular processes, including proliferation and cytoskeletal organisation. We observed that exposure to *B. breve* increases the mRNA abundance of *Mapk14* and carnitine, a metabolites that was found to be altered in the Lz of the BIF group compared to the SPF group, can also promote p38MAPK signalling activation in cardiac tissue [75]. Taken together, our findings reveal that (1) maternal gut microbiota promotes fetal and placental growth in mice and (2) *B. breve* UCC2003 treatment may link to the altered metabolites/nutrient milieu in the mother, affecting placental nutrient transporter abundance and placental barrier thickness for exchange, with effects on fetal growth and development (when compared to GF).

### Limitations of the study

While our study has clear strengths and strong translational implications for pregnancy complication treatments, it has limitations that are important to consider as they impact on the conclusions drawn. First, our study only addresses the effects of *B. breve* UCC2003 in a completely clean and naïve microbiome system (GF model). This is not representative of the human gut environment, and therefore future experiments could include the addition of a SPF group treated with *B. breve* UCC2003 and also a similar group treated with another probiotic species (e.g. *Lactobacillus acidophilus*), or a combination of species. This would help to define *Bifidobacterium*-specific effects (driven by key metabolites), including their efficacy, safety and potential use of probiotics during pregnancy. Moreover, it could be argued that the SPF group interferes in the interpretations of the *B. breve* effects. However, there is lack of fundamental knowledge on what is considered “normal or abnormal” in the GF system, as very little research has been done in understanding the role of the maternal gut microbiota on placental development (SPF vs GF). Therefore, the addition of the SPF group is required to define a baseline for adequate feto-placental growth, and it would also be important to understand how reconstitution of GF mice with SPF microbiota also modulates these responses. As previously mentioned, future work should evaluate the response of pregnant SPF mice to *B. breve* UCC2003 supplementation (using microbial profiling [e.g. shotgun metagenomics] to follow microbiota changes), as well as the efficiency of *B. breve* UCC2003 using other types of mouse models such as antibiotic-treated mice. These animal models may also reduce issues with the immune naïve physiology of the GF system [8]. Unsurprisingly, we did not see a full ‘rescue’ of placental phenotype in the monocolonised GF *B. breve* (BIF) group, compared to the complex microbiota found in SPF dams. However, structural and functional adaptations of these placentas exposed to *B. breve* were adequate enough to ‘rescue’ fetal weight and fetal glycaemia. An array of gut-associated signaling and a diverse metabolite pool are expected to provide more complete placental development. Indeed, other or additional *Bifidobacterium* species and/or strains may be required for placental and fetal development, given strain-specific host physiology responses [42, 76]. Further studies should allow the relative contributions of other microbial- and *Bifidobacterium*-derived factors to be elucidated. Moreover, ideally, future work should analyze fetal and placental growth each day of the supplementation period and use larger cohorts of pregnant mice. It would also be valuable to assess the impact of *B. breve* supplementation from prior to, and/or during the whole pregnancy.

Exploring three different compartments (i.e. mother, placenta and fetus) with respect to metabolites and elucidating their role is a complex process, and makes interpretations and drawing definitive conclusions difficult. Further studies using e.g. 13^C^ labeled *Bifidobacterium* or specific metabolites for tracking experiments may allow more nuanced interactions to be uncovered in future work. Nonetheless, this study has revealed novel roles for the gut microbiota and specifically *Bifidobacterium* and provides the bases for future therapeutic strategies for treating pregnancy complications. These data suggest an opportunity for *in utero* programming through maternal *Bifidobacterium* and associated metabolites. Overall, although our study was performed in mice and is not representative of a clinical scenario, our study highlights the importance of the maternal gut microbiota during gestation and demonstrates that *B. breve* modulates maternal physiology, placental structure and nutrient transporter capacity with impact on fetal glycaemia and fetal growth (Figure 5). Our findings prompt an in-depth investigation into how additional members of the maternal gut microbiota impact on pregnancy outcomes. These future studies are important for the design of novel therapies to combat fetal growth restriction and other pregnancy complications.

**Figure 5.**
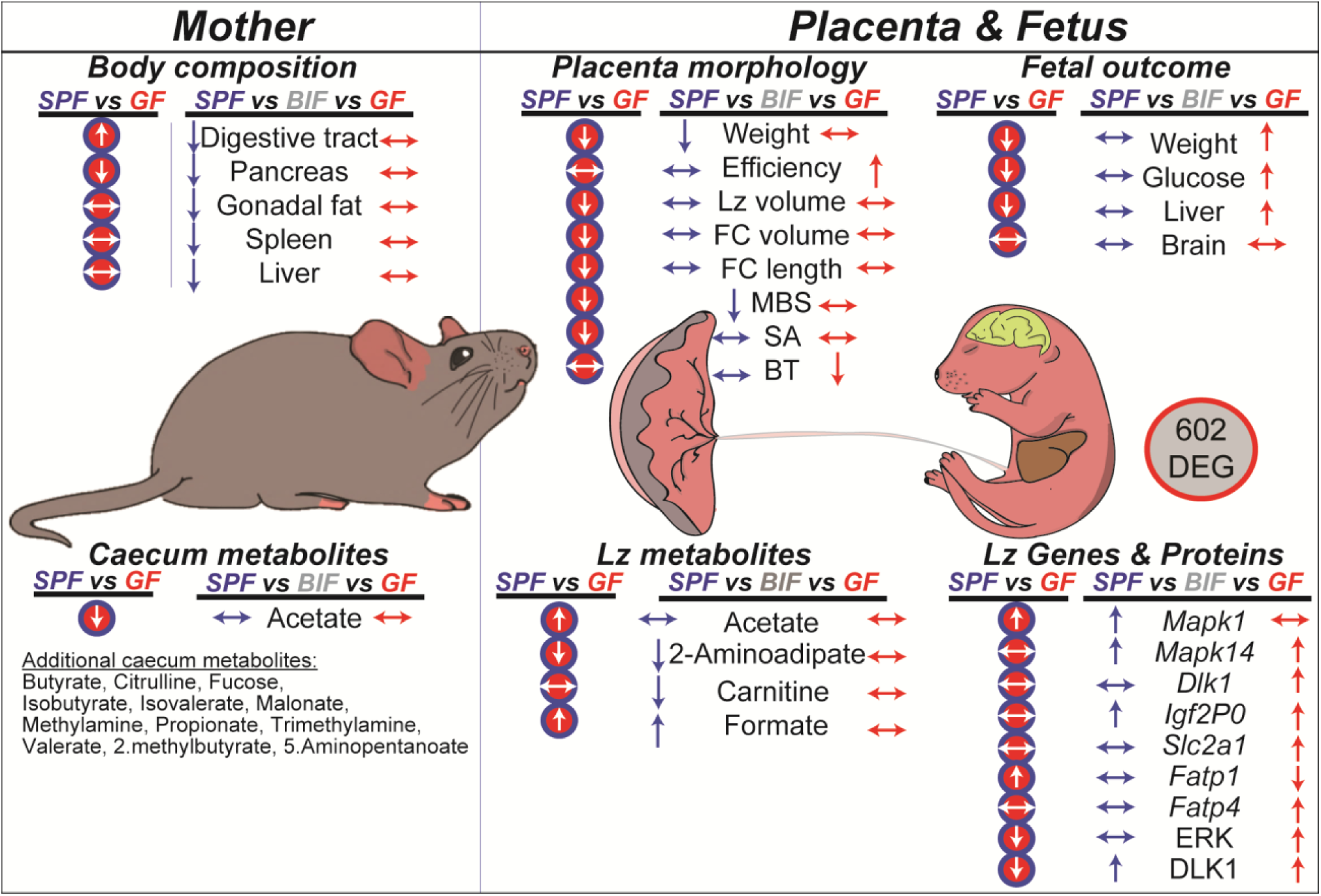
Summary illustration showing the most relevant results on how the maternal gut microbiota and *B. breve* affects mother, placenta and fetus during gestation. The effects of lacking maternal gut microbiota on maternal, placental and fetal phenotype are shown in red circles (SPF vs GF comparisons). Our results suggest that lacking maternal gut microbiota aside from inducing changes in the maternal digestive tract, pancreas and caecum metabolites, has important implications for the correct growth of the fetus and its placenta. The effects of *B. breve* administration compared to the SPF and GF groups are shown in blue and red arrows, respectively. Overall, *B. breve* induces changes in the maternal compartment that affect the structure, metabolome and function of the placenta in association with alterations in fetal metabolism, growth and hepatic transcriptome. Abbreviations: SPF (Specific-Pathogen-Free mouse); GF (Germ-Free mouse); BIF (Germ-Free mouse treated with *B. breve* UCC2003); Lz (labyrinth zone); MBS (maternal blood spaces); FC (fetal capillaries); SA (surface area for exchange); BT (barrier thickness); DEG (differentially expressed genes)

## Ethics approval and consent to participate

All mouse experiments were performed under the UK Regulation of Animals (Scientific Procedures) Act of 1986. The project license PDADA1B0C under which these studies were carried out was approved by the UK Home Office and the UEA Ethical Review Committee.

## Consent for publication

This research was funded in whole, or in part, by the Wellcome Trust [Grant numbers: 220456/Z/20/Z, 100974/C/13/Z and 220876/Z/20/Z]. For the purpose of open access, the author has applied a CC BY public copyright licence to any Author Accepted Manuscript version arising from this submission.

## Author Contributions

JL-T, ZS, ANS-P, LJH designed research; JL-T, ZS, RK, GLG conducted research, JL- T, ZS, RK, MJD contributed analytic tools and performed analysis; DvS contributed reagents; JL-T, ZS, ANS-P, LJH wrote the paper with feedback from all the authors.

## Acknowledgments

Authors would like to thank Dr Ruben Bermejo-Poza (Complutense University of Madrid) for statistical advice and the Ferguson-Smith laboratory (University of Cambridge) for providing the DLK1 antibody.

## Data availability

The fetal liver RNA-Seq raw sequencing data are deposited at the National Center for Biotechnology Information (NCBI) under BioProject PRJNA748000. Relevant data are within the manuscript, individual figures and its Supporting Information files. Additionally, data is available from the corresponding authors on reasonable request. Scripts for differential gene expression analysis can be accessed at GitHub, https://github.com/raymondkiu/Maternal-foetal-microbiota-paper/

**Supplementary Figure 1.**
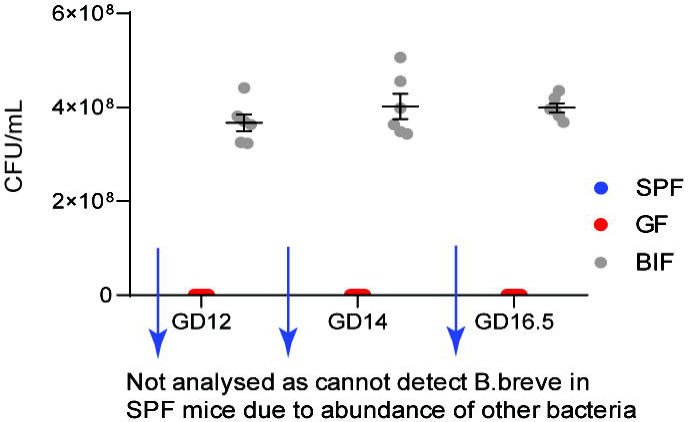
Colonization levels of *B. breve* determined in maternal faecal samples on gestational day (GD), 12 and 14. Analysis performed by two-ways ANOVA (****P<0.0001). Data displayed as mean ± SEM. Number of dams for GF and BIF groups are 5 and 6, respectively. Assessment was performed only on dams sacrificed at GD16.5.

**Supplementary Table 1.**
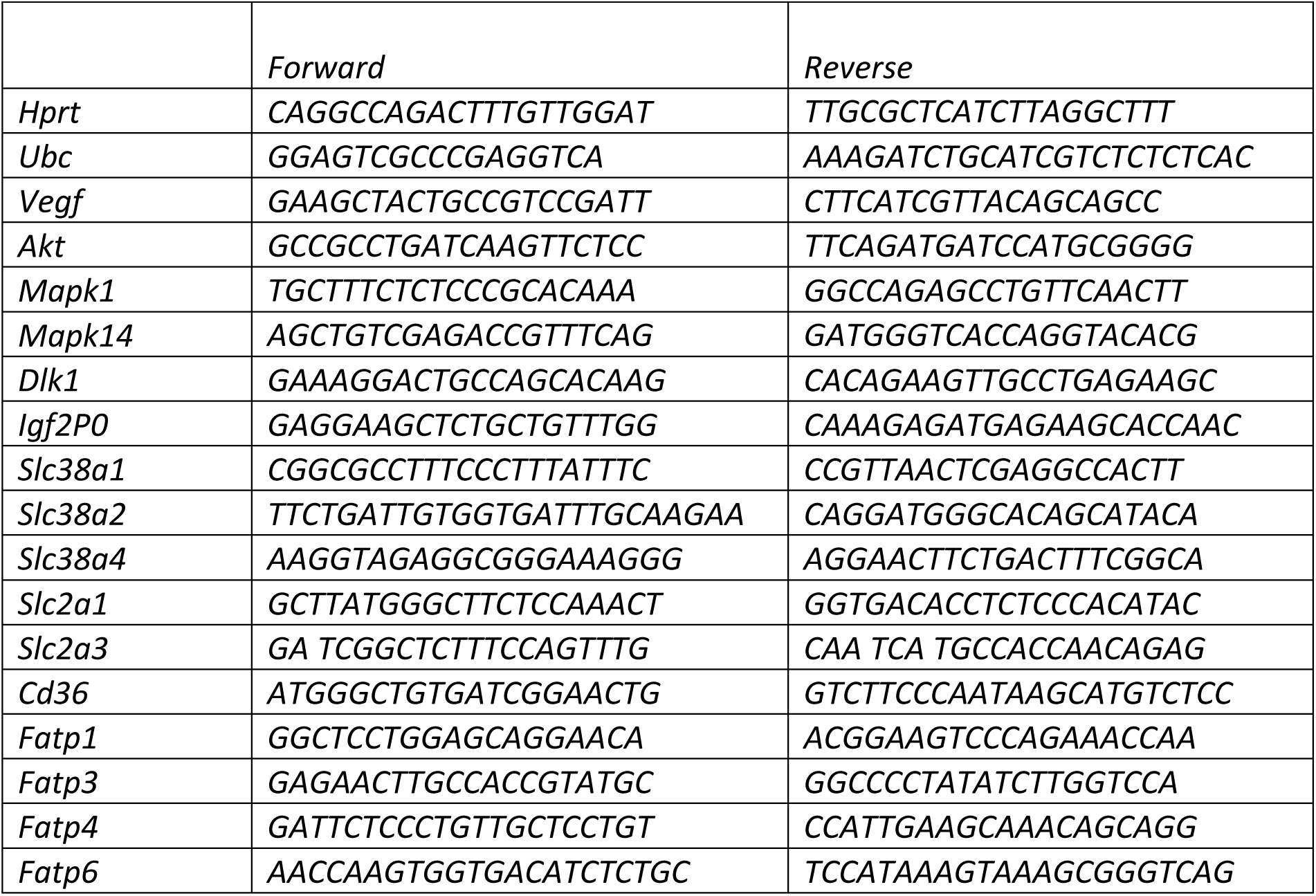
List of primers used for placental labyrinth zone qPCR.

**Supplementary Table 2. List of metabolites analysed in maternal caecum, placental labyrinth zone and fetal liver on day 16.5 of gestation**.

**Supplementary Table 3. List of differentially expressed genes and pathways detected in the liver RNA-Seq on day 18.5 of gestation**.

## References

1. Krajmalnik-Brown R, Ilhan Z-E, Kang D-W, DiBaise JK (2012) Effects of Gut Microbes on Nutrient Absorption and Energy Regulation. Nutr Clin Pract 27:201–214. https://doi.org/10.1177/0884533611436116

2. McDonald B, McCoy KD (2019) Maternal microbiota in pregnancy and early life. Science 365:984–985. https://doi.org/10.1126/science.aay0618

3. Agüero MG de, Ganal-Vonarburg SC, Fuhrer T, et al (2016) The maternal microbiota drives early postnatal innate immune development. Science 351:1296–1302. https://doi.org/10.1126/science.aad2571

4. Koren O, Goodrich JK, Cullender TC, et al (2012) Host remodeling of the gut microbiome and metabolic changes during pregnancy. Cell 150:470–480. https://doi.org/10.1016/j.cell.2012.07.008

5. Nuriel-Ohayon M, Neuman H, Ziv O, et al (2019) Progesterone Increases Bifidobacterium Relative Abundance during Late Pregnancy. Cell Rep 27:730-736.e3. https://doi.org/10.1016/j.celrep.2019.03.075

6. Napso T, Yong H, Lopez-Tello J, Sferruzzi-Perri AN (2018) The role of placental hormones in mediating maternal adaptations to support pregnancy and lactation. Front Physiol 9:. https://doi.org/10.3389/fphys.2018.01091

7. Lamousé-Smith ES, Tzeng A, Starnbach MN (2011) The intestinal flora is required to support antibody responses to systemic immunization in infant and germ free mice. PLoS One 6:e27662. https://doi.org/10.1371/journal.pone.0027662

8. Kennedy EA, King KY, Baldridge MT (2018) Mouse Microbiota Models: Comparing Germ-Free Mice and Antibiotics Treatment as Tools for Modifying Gut Bacteria. Frontiers in Physiology 9:1534. https://doi.org/10.3389/fphys.2018.01534

9. Lu J, Synowiec S, Lu L, et al (2018) Microbiota influence the development of the brain and behaviors in C57BL/6J mice. PLoS One 13:e0201829. https://doi.org/10.1371/journal.pone.0201829

10. Martin AM, Yabut JM, Choo JM, et al (2019) The gut microbiome regulates host glucose homeostasis via peripheral serotonin. PNAS 116:19802–19804. https://doi.org/10.1073/pnas.1909311116

11. Faas MM, Liu Y, Borghuis T, et al (2019) Microbiota Induced Changes in the Immune Response in Pregnant Mice. Front Immunol 10:2976. https://doi.org/10.3389/fimmu.2019.02976

12. Pokusaeva K, Fitzgerald GF, van Sinderen D (2011) Carbohydrate metabolism in Bifidobacteria. Genes Nutr 6:285–306. https://doi.org/10.1007/s12263-010-0206-6

13. Turroni F, Ventura M, Buttó LF, et al (2014) Molecular dialogue between the human gut microbiota and the host: a Lactobacillus and Bifidobacterium perspective. Cell Mol Life Sci 71:183–203. https://doi.org/10.1007/s00018-013-1318-0

14. Turroni F, Milani C, Duranti S, et al (2018) Bifidobacteria and the infant gut: an example of co-evolution and natural selection. Cell Mol Life Sci 75:103–118. https://doi.org/10.1007/s00018-017-2672-0

15. James K, O’Connell Motherway M, Penno C, et al (2018) Bifidobacterium breve UCC2003 Employs Multiple Transcriptional Regulators To Control Metabolism of Particular Human Milk Oligosaccharides. Appl Environ Microbiol 84:e02774–17. https://doi.org/10.1128/AEM.02774-17

16. Food and Agriculture Organization of the United Nations, World Health Organization (2006) Probiotics in food: health and nutritional properties and guidelines for evaluation. Food and Agriculture Organization of the United Nations : World Health Organization, Rome

17. Mazé A, O’Connell-Motherway M, Fitzgerald GF, et al (2007) Identification and Characterization of a Fructose Phosphotransferase System in Bifidobacterium breve UCC2003. Applied and Environmental Microbiology 73:545. https://doi.org/10.1128/AEM.01496-06

18. Cronin M, Akin AR, Collins SA, et al (2012) High Resolution In Vivo Bioluminescent Imaging for the Study of Bacterial Tumour Targeting. PLOS ONE 7:e30940. https://doi.org/10.1371/journal.pone.0030940

19. Hughes KR, Schofield Z, Dalby MJ, et al (2020) The early life microbiota protects neonatal mice from pathological small intestinal epithelial cell shedding. The FASEB Journal 34:7075–7088. https://doi.org/10.1096/fj.202000042R

20. Musial B, Fernandez-Twinn DS, Vaughan OR, et al (2016) Proximity to Delivery Alters Insulin Sensitivity and Glucose Metabolism in Pregnant Mice. Diabetes 65:851–860. https://doi.org/10.2337/db15-1531

21. De Clercq K, Lopez-Tello J, Vriens J, Sferruzzi-Perri AN (2020) Double-label immunohistochemistry to assess labyrinth structure of the mouse placenta with stereology. Placenta 94:44–47. https://doi.org/10.1016/j.placenta.2020.03.014

22. Sferruzzi-Perri AN, López-Tello J, Fowden AL, Constancia M (2016) Maternal and fetal genomes interplay through phosphoinositol 3-kinase(PI3K)-p110α signaling to modify placental resource allocation. Proceedings of the National Academy of Sciences 113:11255–11260. https://doi.org/10.1073/pnas.1602012113

23. López-Tello J, Pérez-García V, Khaira J, et al (2019) Fetal and trophoblast PI3K p110α have distinct roles in regulating resource supply to the growing fetus in mice. Elife 8:. https://doi.org/10.7554/eLife.45282

24. Coan PM, Ferguson-Smith AC, Burton GJ (2004) Developmental dynamics of the definitive mouse placenta assessed by stereology. Biol Reprod 70:1806–1813. https://doi.org/10.1095/biolreprod.103.024166

25. Salazar-Petres E, Carvalho DP, Lopez-Tello J, Sferruzzi-Perri AN (2021) Placental mitochondrial function, nutrient transporters, metabolic signalling and steroid metabolism relate to fetal size and sex in mice

26. Romero-Calvo I, Ocón B, Martínez-Moya P, et al (2010) Reversible Ponceau staining as a loading control alternative to actin in Western blots. Anal Biochem 401:318–320. https://doi.org/10.1016/j.ab.2010.02.036

27. Livak KJ, Schmittgen TD (2001) Analysis of relative gene expression data using real-time quantitative PCR and the 2(-Delta Delta C(T)) Method. Methods 25:402–408. https://doi.org/10.1006/meth.2001.1262

28. Kiu R, Treveil A, Harnisch LC, et al (2020) Bifidobacterium breve UCC2003 Induces a Distinct Global Transcriptomic Program in Neonatal Murine Intestinal Epithelial Cells. iScience 23:101336. https://doi.org/10.1016/j.isci.2020.101336

29. Chen S, Zhou Y, Chen Y, Gu J (2018) fastp: an ultra-fast all-in-one FASTQ preprocessor. Bioinformatics 34:i884–i890. https://doi.org/10.1093/bioinformatics/bty560

30. Kopylova E, Noé L, Touzet H (2012) SortMeRNA: fast and accurate filtering of ribosomal RNAs in metatranscriptomic data. Bioinformatics 28:3211–3217. https://doi.org/10.1093/bioinformatics/bts611

31. Bray NL, Pimentel H, Melsted P, Pachter L (2016) Near-optimal probabilistic RNA-seq quantification. Nat Biotechnol 34:525–527. https://doi.org/10.1038/nbt.3519

32. Zerbino DR, Achuthan P, Akanni W, et al (2018) Ensembl 2018. Nucleic Acids Res 46:D754–D761. https://doi.org/10.1093/nar/gkx1098

33. Pimentel H, Bray NL, Puente S, et al (2017) Differential analysis of RNA-seq incorporating quantification uncertainty. Nat Methods 14:687–690. https://doi.org/10.1038/nmeth.4324

34. Kinsella RJ, Kähäri A, Haider S, et al (2011) Ensembl BioMarts: a hub for data retrieval across taxonomic space. Database (Oxford) 2011:bar030. https://doi.org/10.1093/database/bar030

35. Raudvere U, Kolberg L, Kuzmin I, et al (2019) g:Profiler: a web server for functional enrichment analysis and conversions of gene lists (2019 update). Nucleic Acids Res 47:W191–W198. https://doi.org/10.1093/nar/gkz369

36. Le Gall G (2015) Sample collection and preparation of biofluids and extracts for NMR spectroscopy. Methods Mol Biol 1277:15–28. https://doi.org/10.1007/978-1-4939-2377-9_2

37. Wu J, An Y, Yao J, et al (2010) An optimised sample preparation method for NMR-based faecal metabonomic analysis. Analyst 135:1023–1030. https://doi.org/10.1039/b927543f

38. Lazic SE, Essioux L (2013) Improving basic and translational science by accounting for litter-to-litter variation in animal models. BMC Neuroscience 14:37. https://doi.org/10.1186/1471-2202-14-37

39. Fanning S, Hall LJ, Cronin M, et al (2012) Bifidobacterial surface-exopolysaccharide facilitates commensal-host interaction through immune modulation and pathogen protection. PNAS 109:2108–2113. https://doi.org/10.1073/pnas.1115621109

40. Single-cell RNA-Seq analysis reveals dynamic trajectories during mouse liver development -PubMed. https://pubmed.ncbi.nlm.nih.gov/29202695/. Accessed 27 Jan 2022

41. Dalby MJ, Hall LJ (2020) Recent advances in understanding the neonatal microbiome. F1000Res 9:F1000 Faculty Rev-422. https://doi.org/10.12688/f1000research.22355.1

42. Hughes KR, Harnisch LC, Alcon-Giner C, et al (2017) Bifidobacterium breve reduces apoptotic epithelial cell shedding in an exopolysaccharide and MyD88-dependent manner. Open Biol 7:160155. https://doi.org/10.1098/rsob.160155

43. Miao T, Yu Y, Sun J, et al (2021) Decrease in abundance of bacteria of the genus Bifidobacterium in gut microbiota may be related to pre-eclampsia progression in women from East China. Food & Nutrition Research. https://doi.org/10.29219/fnr.v65.5781

44. Vuong HE, Pronovost GN, Williams DW, et al (2020) The maternal microbiome modulates fetal neurodevelopment in mice. Nature 586:281–286. https://doi.org/10.1038/s41586-020-2745-3

45. Godfrey KM, Haugen G, Kiserud T, et al (2012) Fetal Liver Blood Flow Distribution: Role in Human Developmental Strategy to Prioritize Fat Deposition versus Brain Development. PLOS ONE 7:e41759. https://doi.org/10.1371/journal.pone.0041759

46. López-Tello J, Arias-Álvarez M, Jiménez-Martínez M-Á, et al (2017) The effects of sildenafil citrate on feto-placental development and haemodynamics in a rabbit model of intrauterine growth restriction. Reprod Fertil Dev 29:1239–1248. https://doi.org/10.1071/RD15330

47. López-Tello J, Arias-Alvarez M, Jimenez-Martinez MA, et al (2017) Competition for Materno-Fetal Resource Partitioning in a Rabbit Model of Undernourished Pregnancy. PLOS ONE 12:e0169194. https://doi.org/10.1371/journal.pone.0169194

48. Giussani DA (2016) The fetal brain sparing response to hypoxia: physiological mechanisms. J Physiol (Lond) 594:1215–1230. https://doi.org/10.1113/JP271099

49. Kikuchi K, Ben Othman M, Sakamoto K (2018) Sterilized bifidobacteria suppressed fat accumulation and blood glucose level. Biochem Biophys Res Commun 501:1041–1047. https://doi.org/10.1016/j.bbrc.2018.05.105

50. Wu H, Esteve E, Tremaroli V, et al (2017) Metformin alters the gut microbiome of individuals with treatment-naive type 2 diabetes, contributing to the therapeutic effects of the drug. Nat Med 23:850–858. https://doi.org/10.1038/nm.4345

51. Solito A, Bozzi Cionci N, Calgaro M, et al (2021) Supplementation with Bifidobacterium breve BR03 and B632 strains improved insulin sensitivity in children and adolescents with obesity in a cross-over, randomized double-blind placebo-controlled trial. Clinical Nutrition 40:4585–4594. https://doi.org/10.1016/j.clnu.2021.06.002

52. Sferruzzi-Perri AN, Higgins JS, Vaughan OR, et al (2019) Placental mitochondria adapt developmentally and in response to hypoxia to support fetal growth. Proc Natl Acad Sci USA 116:1621–1626. https://doi.org/10.1073/pnas.1816056116

53. Fuller M, Priyadarshini M, Gibbons SM, et al (2015) The short-chain fatty acid receptor, FFA2, contributes to gestational glucose homeostasis. American Journal of Physiology-Endocrinology and Metabolism 309:E840–E851. https://doi.org/10.1152/ajpendo.00171.2015

54. Aoki R, Kamikado K, Suda W, et al (2017) A proliferative probiotic Bifidobacterium strain in the gut ameliorates progression of metabolic disorders via microbiota modulation and acetate elevation. Sci Rep 7:43522. https://doi.org/10.1038/srep43522

55. Fukuda S, Toh H, Hase K, et al (2011) Bifidobacteria can protect from enteropathogenic infection through production of acetate. Nature 469:543–547. https://doi.org/10.1038/nature09646

56. González Hernández MA, Canfora EE, Jocken JWE, Blaak EE (2019) The Short-Chain Fatty Acid Acetate in Body Weight Control and Insulin Sensitivity. Nutrients 11:1943. https://doi.org/10.3390/nu11081943

57. Hu M, Eviston D, Hsu P, et al (2019) Decreased maternal serum acetate and impaired fetal thymic and regulatory T cell development in preeclampsia. Nat Commun 10:3031. https://doi.org/10.1038/s41467-019-10703-1

58. Priyadarshini M, Thomas A, Reisetter AC, et al (2014) Maternal short-chain fatty acids are associated with metabolic parameters in mothers and newborns. Transl Res 164:153–157. https://doi.org/10.1016/j.trsl.2014.01.012

59. Thorburn AN, McKenzie CI, Shen S, et al (2015) Evidence that asthma is a developmental origin disease influenced by maternal diet and bacterial metabolites. Nat Commun 6:7320. https://doi.org/10.1038/ncomms8320

60. Kim CH (2021) Control of lymphocyte functions by gut microbiota-derived short-chain fatty acids. Cell Mol Immunol 18:1161–1171. https://doi.org/10.1038/s41423-020-00625-0

61. Placental-specific insulin-like growth factor 2 (Igf2) regulates the diffusional exchange characteristics of the mouse placenta | PNAS. https://www.pnas.org/doi/10.1073/pnas.0402508101. Accessed 8 May 2022

62. Constância M, Hemberger M, Hughes J, et al (2002) Placental-specific IGF-II is a major modulator of placental and fetal growth. Nature 417:945–948. https://doi.org/10.1038/nature00819

63. Baumann MU, Deborde S, Illsley NP (2002) Placental glucose transfer and fetal growth. Endocrine 19:13–22. https://doi.org/10.1385/ENDO:19:1:13

64. Dutta-Roy AK (2000) Transport mechanisms for long-chain polyunsaturated fatty acids in the human placenta. Am J Clin Nutr 71:315S–22S. https://doi.org/10.1093/ajcn/71.1.315s

65. Larqué E, Demmelmair H, Klingler M, et al (2006) Expression pattern of fatty acid transport protein-1 (FATP-1), FATP-4 and heart-fatty acid binding protein (H-FABP) genes in human term placenta. Early Human Development 82:697–701. https://doi.org/10.1016/j.earlhumdev.2006.02.001

66. Sebastián D, Guitart M, García-Martínez C, et al (2009) Novel role of FATP1 in mitochondrial fatty acid oxidation in skeletal muscle cells. J Lipid Res 50:1789–1799. https://doi.org/10.1194/jlr.M800535-JLR200

67. Frost JM, Moore GE (2010) The Importance of Imprinting in the Human Placenta. PLOS Genetics 6:e1001015. https://doi.org/10.1371/journal.pgen.1001015

68. Sferruzzi-Perri AN, Sandovici I, Constancia M, Fowden AL (2017) Placental phenotype and the insulin-like growth factors: resource allocation to fetal growth. J Physiol 595:5057–5093. https://doi.org/10.1113/JP273330

69. Yevtodiyenko A, Schmidt JV (2006) Dlk1 expression marks developing endothelium and sites of branching morphogenesis in the mouse embryo and placenta. Developmental Dynamics 235:1115–1123. https://doi.org/10.1002/dvdy.20705

70. Huang C-C, Kuo H-M, Wu P-C, et al (2018) Soluble delta-like 1 homolog (DLK1) stimulates angiogenesis through Notch1/Akt/eNOS signaling in endothelial cells. Angiogenesis 21:299–312. https://doi.org/10.1007/s10456-018-9596-7

71. Forbes K, Westwood M, Baker PN, Aplin JD (2008) Insulin-like growth factor I and II regulate the life cycle of trophoblast in the developing human placenta. American Journal of Physiology-Cell Physiology 294:C1313–C1322. https://doi.org/10.1152/ajpcell.00035.2008

72. Sandovici I, Georgopoulou A, Pérez-García V, et al (2021) The Imprinted Igf2-Igf2r Axis is Critical for Matching Placental Microvasculature Expansion to Fetal Growth

73. Mudgett JS, Ding J, Guh-Siesel L, et al (2000) Essential role for p38α mitogen-activated protein kinase in placental angiogenesis. PNAS 97:10454–10459. https://doi.org/10.1073/pnas.180316397

74. Cuenda A, Rousseau S (2007) p38 MAP-Kinases pathway regulation, function and role in human diseases. Biochimica et Biophysica Acta (BBA) - Molecular Cell Research 1773:1358–1375. https://doi.org/10.1016/j.bbamcr.2007.03.010

75. Fan Z, Han Y, Ye Y, et al (2017) l-carnitine preserves cardiac function by activating p38 MAPK/Nrf2 signalling in hearts exposed to irradiation. European Journal of Pharmacology 804:7–12. https://doi.org/10.1016/j.ejphar.2017.04.003

76. Ruiz L, Delgado S, Ruas-Madiedo P, et al (2017) Bifidobacteria and Their Molecular Communication with the Immune System. Front Microbiol 8:2345. https://doi.org/10.3389/fmicb.2017.02345

