## Supplementary Table 1 for "Maternal gut microbiota *Bifidobacterium* promotes placental morphogenesis, nutrient transport and fetal growth in mice"

| *Supplementary Table 3* | *Forward* | *Reverse* |
| --- | --- | --- |
| *Hprt* | *CAGGCCAGACTTTGTTGGAT* | *TTGCGCTCATCTTAGGCTTT* |
| *Ubc* | *GGAGTCGCCCGAGGTCA* | *AAAGATCTGCATCGTCTCTCTCAC* |
| *Vegf* | *GAAGCTACTGCCGTCCGATT* | *CTTCATCGTTACAGCAGCC* |
| *Akt* | *GCCGCCTGATCAAGTTCTCC* | *TTCAGATGATCCATGCGGGG* |
| *Mapk1* | *TGCTTTCTCTCCCGCACAAA* | *GGCCAGAGCCTGTTCAACTT* |
| *Mapk14* | *AGCTGTCGAGACCGTTTCAG* | *GATGGGTCACCAGGTACACG* |
| *Dlk1* | *GAAAGGACTGCCAGCACAAG* | *CACAGAAGTTGCCTGAGAAGC* |
| *Igf2P0* | *GAGGAAGCTCTGCTGTTTGG* | *CAAAGAGATGAGAAGCACCAAC* |
| *Slc38a1* | *CGGCGCCTTTCCCTTTATTTC* | *CCGTTAACTCGAGGCCACTT* |
| *Slc38a2* | *TTCTGATTGTGGTGATTTGCAAGAA* | *CAGGATGGGCACAGCATACA* |
| *Slc38a4* | *AAGGTAGAGGCGGGAAAGGG* | *AGGAACTTCTGACTTTCGGCA* |
| *Slc2a1* | *GCTTATGGGCTTCTCCAAACT* | *GGTGACACCTCTCCCACATAC* |
| *Slc2a3* | *GA TCGGCTCTTTCCAGTTTG* | *CAA TCA TGCCACCAACAGAG* |
| *Cd36* | *ATGGGCTGTGATCGGAACTG* | *GTCTTCCCAATAAGCATGTCTCC* |
| *Fatp1* | *GGCTCCTGGAGCAGGAACA* | *ACGGAAGTCCCAGAAACCAA* |
| *Fatp3* | *GAGAACTTGCCACCGTATGC* | *GGCCCCTATATCTTGGTCCA* |
| *Fatp4* | *GATTCTCCCTGTTGCTCCTGT* | *CCATTGAAGCAAACAGCAGG* |
| *Fatp6* | *AACCAAGTGGTGACATCTCTGC* | *TCCATAAAGTAAAGCGGGTCAG* |
